# Background check: Mutational input to size variation depends on ancestor’s breeding value

**DOI:** 10.64898/2026.04.01.715985

**Authors:** Lachlan J. King, Katrina McGuigan

## Abstract

The phenotypic effects of mutations often depend on the genetic background, yet general patterns remain poorly resolved. Here, we tested whether genotypes drawn from the same natural population, but differing in their breeding values for a polygenic trait, differed in their contribution of new mutational variation to that trait. We established >200 mutation-accumulation (MA) lines from four *Drosophila serrata* genotypes. Analysing >44,000 wing-size measurements, collected over 30 generations, we quantified mutational variance and mutational bias for size. Genotypes with the smallest and largest breeding values for size contributed similar (statistically indistinguishable) amounts of mutational variance. In contrast, the genotype with an intermediate breeding value exhibited remarkably low (statistically undetectable) mutational variance, low micro-environmental variance, and high line survival over time, consistent with limited mutational decay in fitness. The three genotypes with detectable mutational input showed declines in mean size over time, indicating a consistent mutational bias toward smaller size, as reported in other taxa. The magnitude of this bias appeared genotype dependent, with the MA populations founded from the larger ancestors declining nearly twice as fast as that founded from the smallest ancestor. Together, these results demonstrate substantial heterogeneity in mutational properties among genotypes within a single natural population where the trait value spans a relatively narrow range. Such genotype-specific mutational input is expected to shape both the standing genetic variance and the evolutionary trajectory of polygenic traits.

## Introduction

Evolutionary phenomena, including adaptation, extinction risk, and the maintenance of genetic variance, are impacted by the effects of new mutations, and by how those mutational effects vary with environmental and genetic context (Agrawal & Whitlock, 2010; Carter et al., 2005; Charlesworth, 1990; Hermisson & Wagner, 2004; Jones et al., 2014; Lynch et al., 1999; Walsh & Lynch, 2018; Zeng & Cockerham, 1993). Much of our empirical insights into the nature of context dependent mutational effects comes from manipulative experiments contrasting diverged genotypes with marked fitness differences (e.g., Johnson et al., 2019; Perfeito et al., 2014; Silander et al., 2007). Yet, for many taxa, mutation-selection-drift processes operate within genetically heterogenous outbred populations, and it is at this level we need to extend our empirical understanding of mutational input. Furthermore, while attention has focused on fitness itself, mutation is also the source of variation in phenotypic traits, and mutation may influence patterns of trait variation and evolution (Carter et al., 2005; Johnson & Barton, 2005; Walsh & Lynch, 2018 Chapter 28; Zhang & Hill, 2008). Here, our goal is to extend understanding of variation in mutational properties by determining whether same-population genotypes that differ in their breeding value for a polygenic trait differ in their contribution of new, mutational, variation to that trait.

Size is influenced by many loci throughout the genome (Kemper et al., 2012; Turner et al., 2011; Yang et al., 2010), and size traits typically have substantial genetic variation (Hansen et al., 2011). Taxa show marked divergence in size, including intra-specific latitudinal clines linked to climate (e.g., Kennington & Hoffmann, 2010). There is widespread evidence of selection favouring larger size (Kingsolver & Diamond, 2011), leading to the puzzling question of what keeps organisms small (Blanckenhorn, 2000). Indirect selection (Kingsolver & Diamond, 2011) and cryptic viability (survival) selection on larger individuals (Blanckenhorn et al., 2011; Bonnet et al., 2017; Janeiro et al., 2025; McGuigan & Blows, 2013) can partly resolve this puzzle. However, mutational bias, and context-dependent mutational effects may also contribute through influences on standing genetic variation.

Mutational distributions skewed toward smaller sizes have been reported for nematodes (Ajie et al., 2005; Azevedo et al., 2002; Ostrow et al., 2007), *Daphnia pulex* (Lynch et al., 1998), *Drosophila melanogaster* (Keightley & Ohnishi, 1998; Santiago et al., 1992; Yang et al., 2001) and for knock-out mutations in mice (Reed et al., 2008). These observations are at odds with a common assumption of some models of genetic variation that, while mutations typically decrease fitness, their pleiotropic effects on non-fitness traits are unbiased, equally likely to increase or decrease trait values (Johnson & Barton, 2005; Keightley & Lynch, 2003; reviewed in Chapter 28 of Walsh & Lynch, 2018). Directional bias of pleiotropic effects can generate directional selection on traits that do not directly influence fitness, and hence can contribute to patterns of trait evolution (or stasis) (Tanaka, 2010; Zhang & Hill, 2008). However, not all investigations have supported directional bias of mutations affecting size (McGuigan & Blows, 2013), while some studies have evidenced bias only in some examined genotypes. For example, Azevedo et al. (2002) reported statistical support for mutationally driven decline in size in one strain of *C. elegans*, but not a second strain, which had a threefold lower rate of decline. How pervasive is directional bias of mutations affecting size, does the extent of bias depend on genetic background, and if so, how?

A range of approaches have highlighted the potential adaptive benefit of deleterious mutations amplifying each other’s deleterious effects (accelerating selective elimination: Kimura & Maruyama, 1966; Kondrashov, 1982) or of beneficial alleles (i.e., high fitness genetic backgrounds) weakening the deleterious effects of mutations (reducing deviation of individuals from the fitness optimum: Wagner et al., 1997). However, there is a lack of empirical consensus for a consistent relationship between fitness and the effect sizes of deleterious mutations (Agrawal & Whitlock, 2010, 2012; de Visser et al., 2011; Johnson et al., 2019; Sanjuán & Elena, 2006). Intriguingly, a recent meta-analysis has suggested that directional bias in epistatic effects might be present for size traits in animals. Bourg et al. (2024) found alleles had smaller effects in genetic backgrounds with higher values of size-related morphological traits, suggesting evolvability to larger sizes may be constrained relative to evolution toward smaller size. Consistently, Azevedo et al. (2002) found faster per-generation mutational decline in size in a *C. elegans* population with smaller size (∼20% less body volume). If individuals with higher breeding values for size dampen the effects of new mutations affecting size, this could contribute both to variation in apparent directional bias of mutations, and in the magnitude of new (mutational) variation in size.

In this study, we compare mutational input to size variation of four *Drosophila serrata* genotypes, all derived from the same collection site in Brisbane, Australia. The genotypes (lines) were sampled from across the range of heritable (wing) size in the *Drosophila serrata* Genome Reference Panel (DsGRP: Reddiex et al., 2018) of inbred lines. There is low genomic relatedness among DsGRP lines (< 0.005: Reddiex et al., 2018), ensuring ancestors were genetically independent samples. The DsGRP lines were established by brother-sister inbreeding from field-collected females (Reddiex et al., 2018), a design that limits the opportunity for the population to adapt to laboratory conditions (Mitchell et al., 2025). From each of the four chosen DsGRP ancestors, we established a panel of daughter lines, and maintained them at small population size (brother-sister mating) for 30 generations, allowing spontaneous mutations to fix (or be lost) under random genetic drift (i.e., a Mutation Accumulation, MA, breeding design: Halligan & Keightley, 2009). In each panel, we estimate the mutational variance (*V_M_*, the per-generation increase in variation due to spontaneous mutations), and the directional bias in mutational effects (1*M*, the per-generation change in mean due to fixation of spontaneous mutations) for (wing) size. We test the null hypotheses that genotypes do not differ in mutational variance or directional biases of effects. Based on evidence of negative directional epistasis from line-cross data (Bourg et al., 2024) and of heterogenous magnitude of directional bias of mutations (Azevedo et al., 2002; Ostrow et al., 2007), we predict *V_M_* and 1*M* will negatively covary with ancestral size.

## Methods

### Mutation accumulation and data collection

Ancestors were chosen based on their breeding value for wing size; wing size is correlated with body size in *Drosophila* (De Moed et al., 1997; Robertson & Reeve, 1952). Wing size (Figure S1) was analysed for both males and females from 72 DsGRP lines. The estimated inter-sex genetic correlation was statistically indistinguishable from 1.0 (Supplementary Methods; Figure S2). The DsGRP lines were ranked by male breeding values (Best Linear Unbiased Predictors obtained from a model excluding females) for size, and five lines chosen, ranging from relatively small through relatively large wing size (Figure S2).

The chosen DsGRP lines each founded a unique panel of ≥60 Mutation Accumulation (MA) lines (Table S1): Panel A (DsGRP63), B (DsGRP20), C (DsGRP120), D (DsGRP101) and E (DsGRP170) (ranked from smallest to largest size; Figure S2). Very low viability (and consequent small sample sizes) meant we were unable to estimate *V*_M_ in Panel B (see Supplementary Methods), and results are presented only for the other four panels. Each MA line was maintained through brother-sister inbreeding for 30 generations following protocols detailed in McGuigan et al. (2011) and Mendel et al. (2024); see also Supplementary Methods. All lines from all panels were maintained synchronously under the same conditions.

Replicate single brother-sister pairs were bred each generation, with one pair contributing all parents of the next generation, while two other pairs (vials) supplied flies (≤4 per sex per vial) for wing measurement. Each generation, ∼50% of lines from each panel were assayed, with each MA line sampled every second generation; all lines were sampled in the terminal generation (∼13 repeated samples per MA line; see Supplementary Methods; Table S2). Size was measured for each fly (see Supplementary Methods) and rescaled (mm x 100) ahead of analysis to ensure sufficient resolution of the variance estimates. Observations >4 SD from their respective panel, sex, and generation mean were excluded from analyses (205 wings, <0.5% of all observations). These outliers were distributed across MA lines, and all MA line means were within 4 SD of their relevant panel mean, suggesting the extreme observations reflected factors other than fixation of large-effect mutations. The final dataset contained 44,156 individuals (Table S1).

### Estimating Mutational Variance

Sampling error, segregating mutations, and micro-environmental effects can all influence estimates of mutational variance (Conradsen et al., 2022a; Lynch et al., 1999). Our design minimised these effects in two ways. First, we estimate among-line variance from samples collected over ∼ four generations, with seven of these time-blocks over the duration of the MA. The increased sample size per line provided by this data pooling improved precision (reduced sampling error) (Klein, 1974; Quinn et al., 2006), while the multigenerational spread of the sampling mitigated transient effects of micro-environment and of segregating mutations (Conradsen et al., 2022a; García-Dorado et al., 2000). Second, we estimated mutational variance from the regression of these, relatively robust, among-line variance estimates on time (generations of inbreeding), further reducing the influence of any factors that may upwardly or downwardly bias the estimate within any given sample (time-block) (McGuigan et al., 2011).

We estimated the among line variance per panel per time-block using REML implemented in SAS software (version 9.4, Copyright 2020, SAS Institute Inc.) to fit the model:

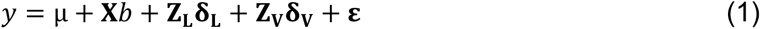

where *y* was the vector of wing size measurements, μ was the intercept (mean size) and 𝐗 was the design matrix relating observations to their level of the categorical fixed-effects (*b*): time-block (accounting for change in mean size over time due either to evolution or environmental trends), micro-environment (accounting for size variation due to uncontrolled heterogeneity in conditions among generations within a time-block: Figure S3), sex, sex by micro-environment interaction, and the researcher recording the landmarks. Mutations had the same (within statistical limits) effect in both sexes (Supplementary Methods), and we therefore estimated the among-line variance estimates from sexes pooled. 𝐙_𝐋_ and 𝐙_𝐕_ were the design matrices relating observations to their random effects of MA line and replicate rearing vial (nested within MA line and generation), respectively. The among line variance 𝛅_𝐋_was modelled using an unstructured covariance matrix to estimate within time-block variance and among time-block covariance. Replicate rearing vial (𝛅_𝐕_), which accounted for little variation in size, was modelled as constant (homogeneous) among micro-environments and time-blocks.

The residual, **ε** (variation among individuals within vials), was modelled as a diagonal matrix with micro-environment-specific variances, allowing us to account for any heterogeneity in residual variance among sampling points (generations).

For each panel, to estimate the mutational variance (𝑉_#_: the per generation increase in phenotypic variance due to mutation), we regressed the seven REML estimates of among-line variance against the panel average number of generations of brother-sister inbreeding at each of these time-blocks (Table S2). To account for any residual segregating genetic variation in the ancestor, and ensure an unbiased estimate of the slope, the intercept was estimated, not constrained to zero (McGuigan et al., 2011). The slope, *b*, was used to estimate: 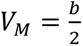 (Lynch & Walsh, 1998, eqn. 12.4a, pp. 331). To test the null hypothesis of no mutational variance (𝑏 = 2𝑉_#_ = 0) we followed a REML-MVN approach (Houle & Meyer, 2015). The “mvrnorm” function in the MASS R package (Ripley et al., 2025; Venables & Ripley, 2002) was used to draw 10,000 samples from the multivariate normal (MVN) distribution, *N* ∼ (θ^0^**, V**), where θ^0^ was the vector of observed (REML) covariance parameter estimates, and **V** was the asymptotic variance-covariance matrix, both derived from model (1). We applied the regression analysis to the 10,000 REML-MVN samples per panel, and accepted the null hypotheses of no mutational variance when the lower 5% REML-MVN confidence interval of the slope included zero.

To compare 𝑉*_M_* to that reported for size traits, we expressed it as mutational heritability, 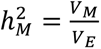 and as the expected proportional change under a unit strength of selection (opportunity for selection, 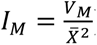) (Garcia-Gonzalez et al., 2012; Houle, 1992). Environmental variance, *V_E_*, was calculated as 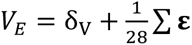, *i.e.*, as the sum of variance among replicate rearing vials (δ_*_) and the average residual variance (**ε**), from model (1). *I_M_* was estimated where mean size, 𝑋^6^, was the experiment-wide panel mean, or the generation 1 (ancestor) mean. Confidence intervals of these scaled parameters were estimated by applying the equations to the REML-MVN samples; for *I_M_*, the REML-MVN samples of *V_M_* were randomly combined with 10,000 samples drawn from a normal distribution defined by the estimate of mean size and its standard deviation.

### Estimating Mutational Bias

Directional bias in mutational effects is evidenced by evolution of panel mean size over time. Size fluctuated markedly among generations, with highly concordant patterns across all panels (Figure S3). Such variation is common in multigenerational laboratory experiments, reflecting minor uncontrollable changes in environmental conditions. For panel C, we had strong evidence of no mutational variance in size (see Results), and therefore designated C as a control population to account for temporally heterogenous micro-environmental effects (sensu Falconer & Mackay, 1996 pg 195). For panels A, D and E, each individual fly’s wing size was expressed as a deviation for the respective generation and sex-specific C panel mean. These deviations were analysed using a maximum likelihood fit (in SAS software version 9.4, Copyright 2020, SAS Institute Inc.) of a mixed model modified from model (1): landmarker and sex were omitted while panel (categorical), generations of inbreeding (continuous) and their interaction were fit as fixed effects. The random effects were as in model (1), but modelled independently for each panel. We applied a Satterthwaite approximation for the degrees of freedom, and tested the null hypothesis that mutational directional bias was the same in all panels. We implemented a planned comparison to test for shared slopes among each pair of panels, and finally whether each panel had a non-zero slope.

The panel-specific slopes were retained as estimates of the directional mutational bias, Δ𝑀, and are reported both on the measurement scale (mm/generation) and relative to the mean size (% of change in mean size per generation). Confidence intervals were estimated from 10,000 random samples drawn from the normal distribution defined by the slope estimate (Δ𝑀) and model-estimate standard deviation of that slope, combined with the set of 10,000 sample estimates of mean size (detailed above). While non-linear evolution of trait mean occurs when the magnitude of mutational effect coevolves with the trait value, short-term experiments such as this are not expected to capture this non-linear signal (Halligan & Keightley, 2009). Visual inspection of the data did not suggest non-linear (accelerating or decelerating) change in mean size with a panel.

## Results

Mutational variance (*V_M_*) for size followed an intriguing U-shaped distribution across the ancestral size distribution, with substantially higher *V_M_* for panels founded by relatively small (A) or large (E) ancestors (Figure 1A; Table 1). Non-zero mutational variance was strongly supported for all panels except C, founded by the intermediate-sized ancestor (Figure 1A; Table 1). Panel C also had relatively little micro-environmental variance (*V*_E_), ∼60% lower than that observed for the relatively extreme genotypes (panels A and E: Figure 1B; Table 1).

**Figure 1.**
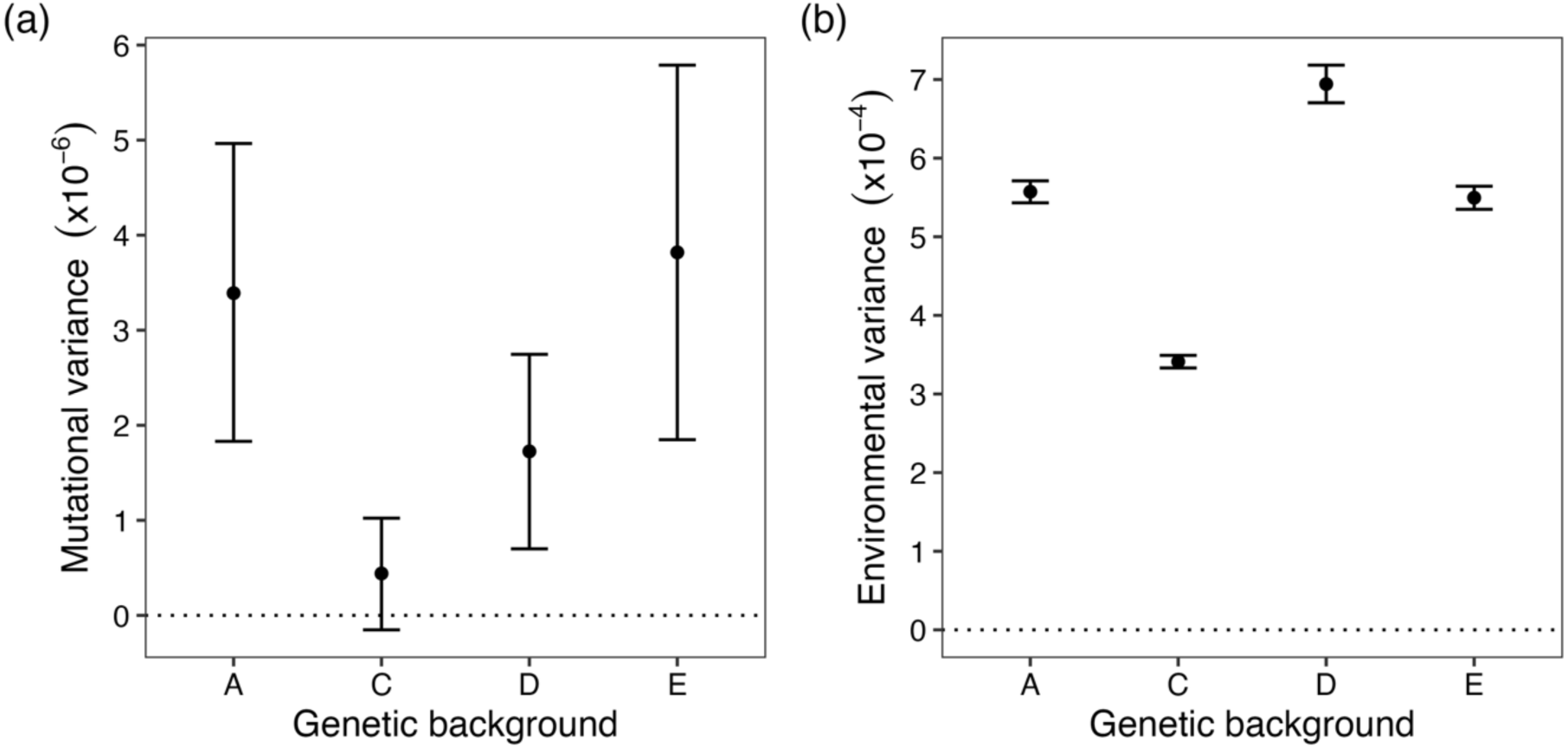
Mutational (a) and environmental (b) variance of wing size in four genetic backgrounds. Points represent estimates and bars the 90% REML-MVN confidence intervals. a) V_-_, estimated from regression of among-line variance on generation. b) V., the sum of among-vial variance and the average (across sampling generations) within vial (residual) variance. In both panels, the dotted line indicates zero, where the null hypothesis of zero variance was accepted when the lower CI was below zero.

**Table 1.** Parameter estimates (and 90% REML-MVN confidence intervals) for each Panel.

| Panel | $V_M$ ($\times 10^{-6}$)<sup>[1]</sup> | $V_E$ ($\times 10^{-4}$)<sup>[2]</sup> | $\bar{X}$ (mm)<sup>[3]</sup> | $h_M^2$ ($\times 10^{-3}$)<sup>[4]</sup> | $I_M$ ($\times 10^{-6}$)<sup>[5]</sup> |
| --- | --- | --- | --- | --- | --- |
| A | 3.90<br>(1.83, 4.96) | 5.57<br>(5.43, 5.71) | 1.31<br>(1.22, 1.41) | 6.08<br>(3.29, 8.90) | 1.98<br>(1.04, 2.98) |
| C | 0.44<br>(-0.15, 1.02) | 3.41<br>(3.33, 3.49) | 1.38<br>(1.28, 1.48) | 1.28<br>(-0.45, 3.00) | 0.23<br>(-0.08, 0.55) |
| D | 1.73<br>(0.70, 2.75) | 6.94<br>(6.70, 7.18) | 1.41<br>(1.30, 1.53) | 2.47<br>(1.00, 3.97) | 0.87<br>(0.35, 1.42) |
| E | 3.82<br>(1.85, 5.79) | 5.50<br>(5.35, 5.64) | 1.43<br>(1.32, 1.54) | 6.95<br>(3.36, 10.53) | 1.88<br>(0.89, 2.93) |

<sup>[1]</sup> Mutational variance, $V_M = \frac{b}{2}$ , where $b$ is the slope of per timeblock among-line variance estimates regressed against the corresponding number of generations of brother-sister inbreeding (Table S2); <sup>[2]</sup> $V_E$ , environmental variance, the sum of the average residual variance and the variance among replicate rearing vials. <sup>[3]</sup> Mean size, $\bar{X}$ , estimated from experiment-wide data per panel (see Table S3 for generation 1 estimates). <sup>[4]</sup> Mutational heritability, $h_M^2 = \frac{V_M}{V_E}$ . <sup>[5]</sup> Mean standardised mutational variance (opportunity for selection), $I_M = \frac{V_M}{\bar{X}^2}$ .

Conclusions about the relative magnitude of *V_M_* (or of genetic variance) depend on the comparison scale (Conradsen et al., 2022a; Hansen et al., 2011), with the mean-standardised scale considered most informative for size traits (Hansen & Houle, 2008; Ostrow et al., 2007). Relative to published estimates for size, our estimates were in the mid-range when considered relative to environmental variance (ℎ^%^ : Figure 2A), while when considered relative to mean size (𝐼_#_), estimates for panels A, D and E were comparable to other *Drosophila* studies, but panel C’s estimate was the smallest recorded (Figure 2B).

**Figure 2.**
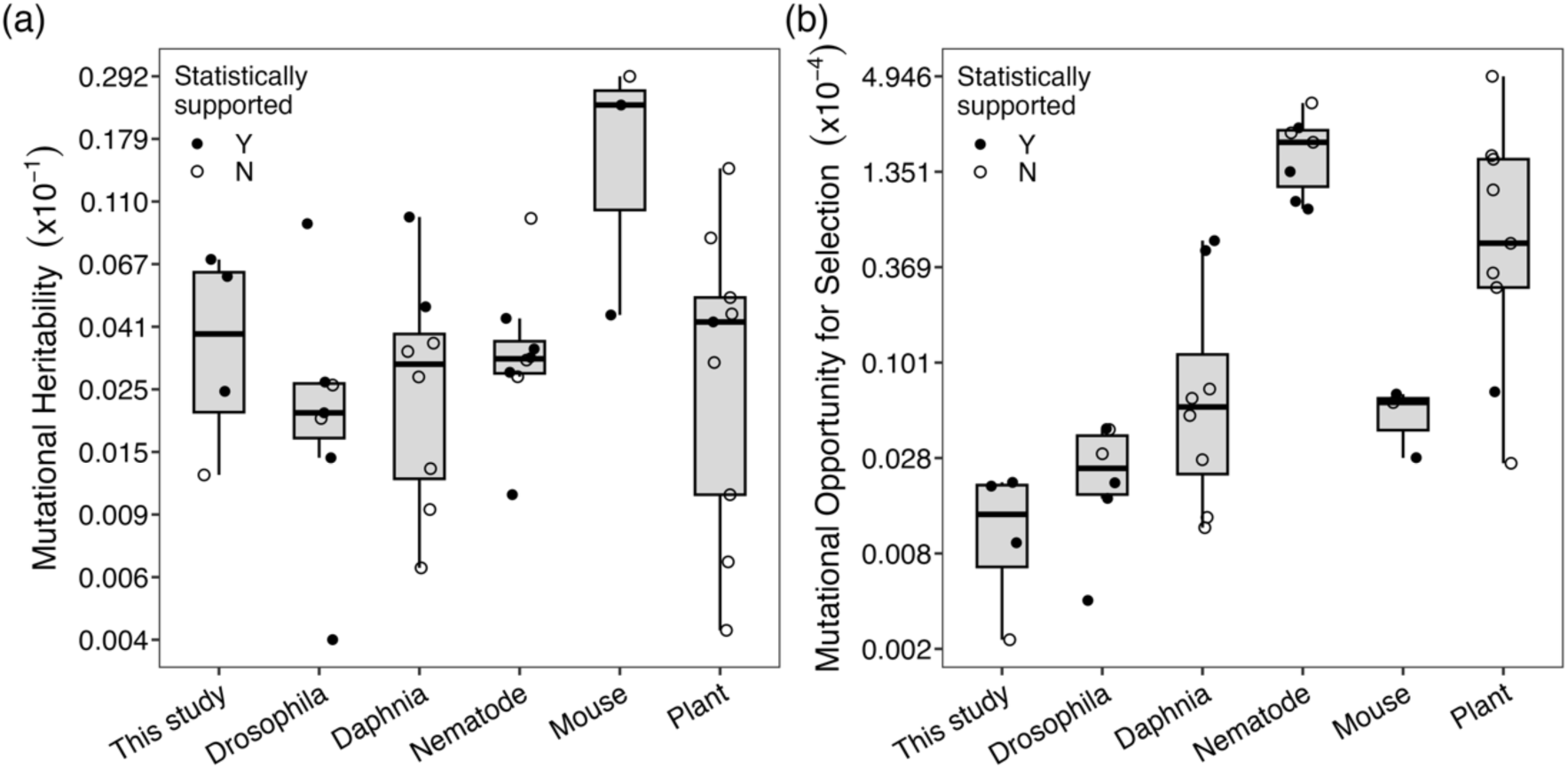
Comparison of mutational variance on A) variance-standardised, ℎ^%^, and B) mean-standardised, 𝐼_#_, scales. Estimates from this study (Table 1) are compared to published estimates for size traits (see Table S1A and C of Conradsen et al., 2022a; Conradsen et al., 2022b). Estimates are plotted on log-scale, but y-axis labels indicate the original (ℎ^%^ or 𝐼_#_) scale. Estimates were considered statistically distinct from zero (Y: closed symbol) if the original authors reported statistical support, or (approximate) standard errors (SE) and the mutational parameter was > 2 SE above zero. Where information was not provided to assess statistical support, estimates were conservatively assumed non-significant. Drosophila: *D. melanogaster* (García-Dorado et al., 2000; Houle & Fierst, 2013; Santiago et al., 1992; Wayne & Mackay, 1998) and *D. serrata* (Dugand et al., 2021); Daphnia: *D. pulex* (Lynch, 1985; Lynch et al., 1998); Nematode: *Caenorhabditis elegans* (Azevedo et al., 2002; Estes et al., 2005; Ostrow et al., 2007), *C. briggsae* and *Oscheius myriophila* (Ostrow et al., 2007); Mouse: *Mus musculus* (Bailey, 1959; Festing, 1973); Plant: *Amsinckia douglasiana* (Schoen, 2005), *Arabidopsis thaliana* (Chang & Shaw, 2003; Kavanaugh & Shaw, 2005) and *Zea mays* (Russell et al., 1963).

The small effective population size of MA lines greatly reduces the efficacy of selection but cannot completely eliminate its influence. Deleterious mutation load can lead to line extinction and, if lines that become extinct had relatively extreme trait values, this line extinction deflates the estimated among-line variance (Lynch et al., 1999; Mackay et al., 1995). Notably, only one panel C MA line became extinct (in the final generation of the experiment), while 7, 9 and 32 lines went extinct from panels A, E and D, respectively (Figure 3; Table S1). Thus, line extinction cannot have contributed to the low *V*_M_ of the C panel, but may have contributed to the lower *V*_M_ in panel D compared with panels A and E. Some MA lines were resurrected by flies emerging from multi-parent vials (∼ 20 parents, with offspring being a mix of full and half siblings and double first cousins: see Supplementary Methods). This transient population expansion could have allowed selection to eliminate some mutations, downwardly biasing the estimate of *V*_M_. Again, panel C was the least and D the most affected by this fitness-related difference in opportunity for selection (2, 8, 15 and 28 C, A, E and D, panel lines experienced this larger population size, respectively: see Supplementary Methods). The relatively low fitness of the D panel was context dependent: most line extinction (and resurrection) occurred after all experimental lines were moved to a different controlled environment facility (Figure 3; Figure S3).

**Figure 3.**
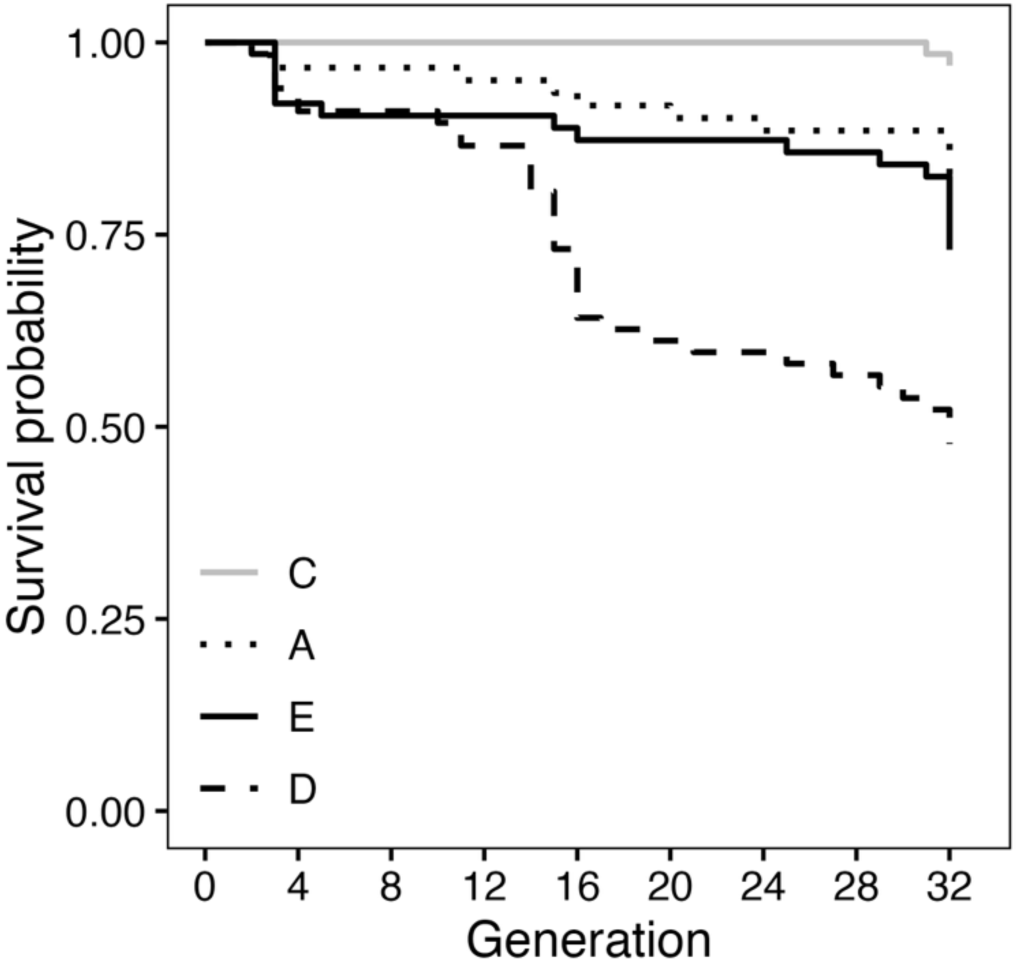
Kaplan-Meier survival curves of MA line survival in each of the four independent mutation accumulation panels. Survival significantly differed among panels (Cox proportional hazard regression model fits for models with panel-specific versus common curve compared using LRT: Ξ^2^ = 50.69, d.f. = 3, p < 0.001). Analyses were conducted using the R package “survival” (Therneau & Grambsch, 2000; Therneau et al., 2024).

### Directional bias of mutational effects

Evolution of mean size as mutations accumulate would indicate either more frequent, or larger average effect of mutations in the evolved direction. Mean size significantly declined in all panels, but ∼twice as fast in panels D and E relative to A (Figure 4; Table S3). We rejected the null hypothesis of equal rates of evolution of mean size among panels (panel x generation interaction: *F*_2,79.7_ = 8.55, *P* = 0.0004), with planned contrasts supporting a difference of A from both D and E, but not between D and E (Figure 4). As a proportion of ancestral (generation 1: Table S3) size, the rate ranged from a 0.021% (A) to 0.044% (E) decline per generation (Figure 4). This was comparable to, but at the slow end, relative to the mutational decline in size in *C. elegans* (estimates from two genetic backgrounds and three independent MA experiments ranged from 0.02% to 0.07% of the ancestor’s mean size: Ajie et al., 2005; Azevedo et al., 2002; Ostrow et al., 2007), and slow to relative to *Oscheius myriophila* (0.09 – 0.14%) and to *C. briggsae* (0.14 – 0.17%) (Ostrow et al., 2007).

**Figure 4.**
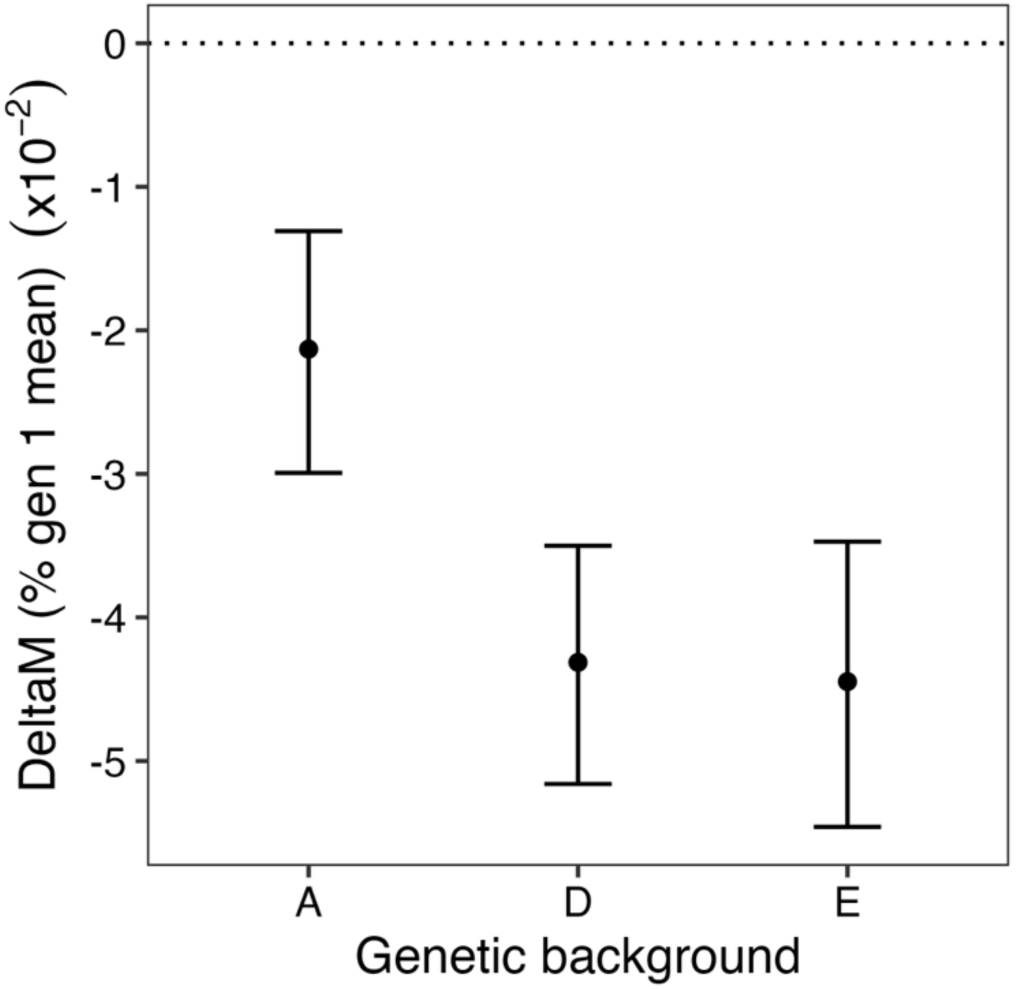
The per generation rate of change in mean size (± 90% CI) under mutation-drift evolution for each of the three evolving panels (A, D and E). Rate is expressed as a percentage of the ancestral mean size (near identical estimates for percent of experiment-wide mean: Table S3). Planned contrasts within the fit model (on the mm scale) supported significantly slower decline in panel A relative to both D (*F*_1,86.3_=12.79, *P* =0.0006) and E (*F*_1,84.5_=11.88, *P* =0.0009), but no difference in this mutational bias between D and E (*F*_1,83,8_=0.06, *P* =0.8010).

## Discussion

Heterogeneity among estimates of mutational characteristics is common. If these differences arise due to correlation of mutational characteristics and trait value, selection and mutation may interact to bias or constrain the direction of evolution (Carter et al., 2005; Tanaka, 2010; Zhang & Hill, 2008). Here, we observed covariation of two mutational characteristics with breeding values for size: i) mutational variance declined toward intermediate breeding values (C) (Table 1; Figures 1 and 2) and; ii) the genetic background with the smallest breeding value (A) exhibited weaker mutational bias toward smaller size (Figure 4). Both patterns suggest the potential for mutation to influence evolutionary patterns.

The most striking results of the current study were the absence of detectable mutational variance and low non-mutational (micro-environmental) variance for size in panel C (Figure 1, Table 1). This pattern indicates a high level of phenotypic robustness—insensitivity of the trait to both genetic and environmental perturbations. Robustness is well documented, and may evolve adaptively to maintain phenotypes near high fitness values (de Visser et al., 2003; Wagner et al., 1997). Although some empirical studies support such a negative relationship between genotypic fitness and the magnitude of mutational or environmental variance (e.g., Baer, 2008; Novella et al., 2013; Sharp & Agrawal, 2012; Sztepanacz et al., 2017; Zheng et al., 2020), others support the opposite pattern (e.g., Butković et al., 2020; Goldstein et al., 2025; Johnson et al., 2019). These conflicting observations may reveal the absence of any universal pattern. Alternatively, given the theoretical importance of local curvature in the adaptive landscape (de Visser et al., 2003; Diaz-Colunga et al., 2023), they may reflect differences in where experimental populations lie on that landscape.

Empirically, mutational variance for size often varies substantially across studies and taxa, but shared designs are rare, and genotypic versus experimental sources of variation are frequently confounded (Conradsen et al., 2022a). Previous within species contrasts (e.g., in *D. melanogaster* and rhabditid nematodes) reported small (<20%; Ostrow et al., 2007) to moderate (∼50%; Houle & Fierst, 2013) differences in mutational variance for size between pairs of genotypes, contrasting with the high (∼80-90%: Table 1) reduction of *V_M_* in C compared to A or E. Here, the difference in ancestral size (ratio of smaller to larger) between C and either A or E was <5% (Table 1), slightly less than between compared pairs of *C. elegans* and *C. briggsae* genotypes (∼5-15%: Ostrow et al., 2007). The nematode genotypes associated with slightly larger size also had slightly higher mutational variance (Ostrow et al., 2007). However, it is unknown where in the size distribution of the population the two compared genotypes were positioned, and thus whether the pattern is consistent with increased mutational variance toward the centre or the upper tail of the size distribution.

The robustness of size (and fitness) to mutational and environmental perturbations of the intermediate size C genotype is suggestive of stabilising selection on size in the natural population from which the DsGRP lines were sampled, and could bias the population toward evolutionary stasis in size. However, this interpretation should be treated cautiously given the lack of empirical support for a consistent relationship between fitness and mutational effects (detailed above), the lack of direct estimation of selection in the wild *D. serrata* population, and fact that inference rests on a single genotype with an intermediate breeding value. Nonetheless, these results, and the evidence of slight positive association between the magnitude of mutational variance and mean size reported for conspecific *Caenorhabditis* genotypes are not consistent with the evidence of Bourg et al. (2024), which implicated decreasing allelic effects in genetic backgrounds conferring larger size.

Persistent directional bias in the effects of new mutations is intriguing as it could generate cryptic evolution (Hadfield et al., 2011; McGuigan et al., 2011) and help to reconcile evidence for pervasive positive selection on size (Kingsolver & Diamond, 2011) with the lack of consistent evolution toward larger size (Gotanda et al., 2015; Merilä et al., 2001). Here, mutation decreased average size, consistent with prior studies across taxa (reviewed in Azevedo et al., 2002; McGuigan & Blows, 2013; Ostrow et al., 2007). The mutational bias toward smaller size was markedly weaker in the genotype with the smallest breeding value (A: −0.02% per generation) than in the genotype with the largest (E: −0.04%) (Figure 4). Evolutionary stasis under positive selection on size would be most strongly supported by just such a stronger negative mutational bias in individuals carrying more size increasing alleles (here, E). However, we are again cautious about such an interpretation of our results. Published estimates of mutational bias vary even for the same genotype. For example, for body volume in the N2 strain of *C. elegans*, estimates ranged from −0.02% to −0.06% per generation across experiments on the same mutation accumulation lines, and among independently replicated MA panels, with only two of three estimates statistically distinct from zero (Ajie et al., 2005; Azevedo et al., 2002). Thus, the difference observed here may reflect chance sampling effects from the same underlying distribution of mutational effects. Nonetheless, the observed positive correlation between heritable size and mutational bias is intriguing and warrants further investigation.

### Conclusions and future directions

Mutation both defines the phenotypic values readily accessible to future evolution, and reflects the action of past evolutionary processes (do O & Whitlock, 2023; Jones et al., 2014; Svensson & Berger, 2019), yet its properties remain poorly understood. This study revealed several intriguing patterns, including marked mutational (and environmental) robustness of a genotype near the population mean size, and reduced bias in mutational effects toward smaller sizes in a genotype from the lower tail of the size distribution. Assessing the generality of these results will require additional estimates of mutational effects, particularly replication among genotypes with similar breeding values and extension to other polygenic traits. However, precise estimation of quantitative mutational parameters depends on large, costly experiments, posing a substantial barrier to progress. The growing availability of large pedigrees from natural populations with phenotypic information may therefore present an underutilised resource for linking trait breeding values with mutational effects (Dugand et al., 2021; McGuigan et al., 2015).

## Data and code availability

All data and scripts are available from King and McGuigan (2026).

## Author contributions

LJK and KM designed research; LJK performed research; LJK and KM analysed data and wrote the paper.

## Supporting information

Supplementary methods and results

## Acknowledgements

Nick Appleton and Derek Sun were instrumental in managing the large number of lines over the 18 months this experiment ran. We are also appreciative of the commitment and hard work of Ash Kannan, Alex Ichim, and Cara Conradsen in maintaining and phenotyping the lines, and of Wasana Nidarshani and Indigo Grigg’s contributions to phenotyping. We thank Henrique Teotónio and Greg Walter for feedback on earlier drafts.

