## Supplementary methods and results for "Background check: Mutational input to size variation depends on ancestor’s breeding value"

#### WING SIZE MEASUREMENT

Methods for wing measurement differed slightly between the DsGRP and MA lines. In the DsGRP, one wing per fly (typically the left) was removed and photographed (following protocols detailed in Conradsen et al., 2022). In the MA lines, wings on intact flies were photographed using a "wing machine" (Houle et al., 2003) to hold the wing in position. The same Leica microscope (model MZ8 at 5x magnification) with attached Leica camera (model IC90E) was used to image wings in both datasets. Images were randomised within a generation, and nine landmarks recorded (Figure S1) on each wing image (TPSdig2 version 2.31: Rohlf, 2017) by one of four researchers ("landmarkers"). In the DsGRP experiment, coordinates of all nine recorded landmarks were aligned (MorphoJ version 1.06 Klingenberg, 2011) and centroid size (Klingenberg, 2016; Rohlf, 1999) retained as a measure of overall wing size. In the MA lines, we followed Sztepanacz and Houle (2021) in excluding internal landmarks to estimate centroid size.

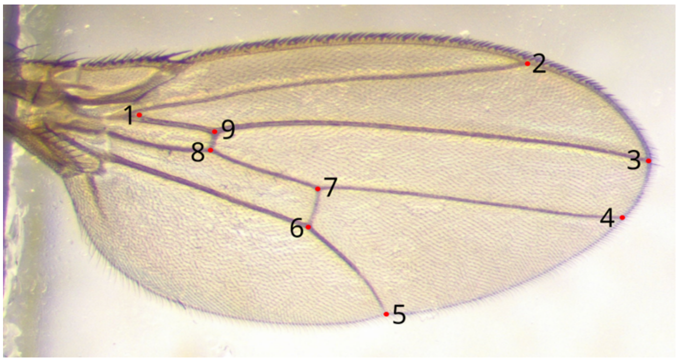


**Figure S1** – Example wing showing the placement of the nine landmarks. All landmarks were analysed to estimate centroid size of DsGRP line flies. For MA line flies, only landmarks 1 through 5 were analysed. Landmarks are defined by the intersection of wing veins and crossveins or wing veins and wing margins.

### ANALYSIS OF DSGRP WINGS

For each of 73 DsGRP lines, wings of three flies per sex from each of two independent rearing vials were measured. We fit a mixed model to data from both sexes using restricted maximum likelihood (REML) implemented in SAS (version 9.4, Copyright 2020, SAS Institute Inc.):

$$y = \mu + \text{Sex} + \text{Line} + \text{Vial} + \varepsilon$$

where *y* was the vector of observations of size (rescaled by multiplying by 100 to improve resolution of small magnitude random effects) and μ the population-mean size; Sex was fit as a categorical fixed effect; the random residual (ε) was modelled as a diagonal matrix, estimating variance per sex. The random effects of among-line and among replicate rearing vial (nested within line) were each modelled as an unstructured correlation matrix, estimating the variance per sex and the inter-sex correlation. At the among-line level, we applied log-likelihood ratio tests to the null hypotheses that the estimated inter-sex correlation was: i) 0.0 or ii) 1.0, using the PARMS statement to hold the correlation estimate to 0.00001 or 0.99999, respectively.

The estimated inter-sex genetic correlation was 0.9305 (SE: 0.0573), and we rejected the null hypothesis that it was 0.0 ($X^2 = 44.22$, d.f. = 1, $P < 0.0001$) but failed to reject the null hypothesis that it was 1.0 ($X^2 = 1.49$, d.f. = 1, $P = 0.2228$). The sex-specific breeding values (Best Linear Unbiased Predictors) are plotted in Figure S2: The chosen DsGRP ancestors were not unique – other DsGRP lines with a similar wing size could have been chosen instead. Our goal in selecting ancestors was to sample from across a range of sizes, not to sample specific sizes or lines. The chosen DsGRP lines each founded a unique panel of ≥60 Mutation Accumulation (MA) lines (Table S1).

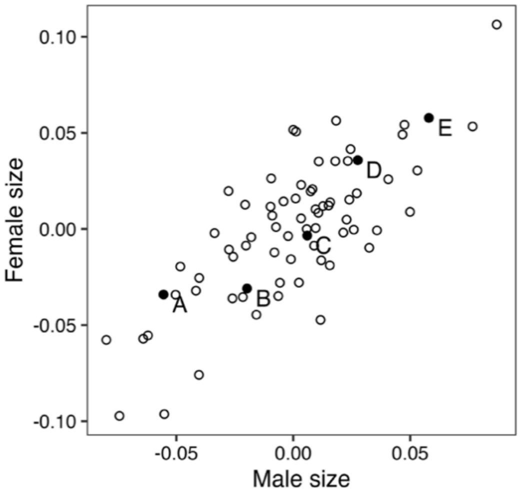


**Figure S2** - The distribution of male and female Best Unbiased Linear Predictors (BLUPs) for wing size in 72 DsGRP lines. BLUPs are plotted as deviations (in units mm) from the sex-specific population (i.e., all DsGRP lines) mean. Filled points correspond to the five chosen MA ancestors: A (DsGRP63), B (DsGRP20), C (DsGRP120), D (DsGRP101) and E (DsGRP170).

**Table S1 -** Experimental sample sizes. The number of MA lines initiated (Gen 1) and surviving to the end of the experiment (Gen 32), and the number (%) of the extant lines sampled for wings in a generation, on average across all generations. Also presented are the number of wings sampled, in total (including outliers that were omitted prior to analysis) and on average per generation (excluding generation 32 when all, rather than only odd or even numbered, lines were sampled). NB: two lines (1xB and 1xC) were lost due to administrative error (i.e., not due to failure to produce live flies).

| | Number of lines | | | Number of wings | |
|---|---|---|---|---|---|
| Panel | Gen 1 | Gen 32 (% survival) | Mean sampled (mean %) | Total | Mean per generation |
| A | 61 | 54 (89%) | 25.7 (45.4%) | 11351 | 392.8 |
| B | 61 | 26 (43%) | 14.3 (34.6%) | 5408 | 192.9 |
| C | 68 | 66 (97%) | 32.3 (47.6%) | 14101 | 489.2 |
| D | 67 | 35 (52%) | 20.6 (40.9%) | 8233 | 296.1 |
| E | 63 | 52 (83%) | 24.7 (44.2%) | 10676 | 367.5 |
| Combined | 320 | 233 (73%) | 23.5 (42.5%) | 49769 | 1738.5 |

### FLY MAINTENANCE AND MUTATION ACCUMULATION LINE INBREEDING

All flies (DsGRP and MA lines) were maintained on standard laboratory media, a mixture of yeast, raw sugar, agar, and antimicrobial agents (Kannan et al., 2023) at 25° C with a 12hr:12hr light:dark cycle, and 14-day generation time. Breeding and virgin holding vials contained 10mL or 5mL of media, respectively. Flies were sexed under $CO_2$ anaesthesia.

For each MA line, 10 replicate single pair rearing vials were set up each generation, all derived from the same parental vial (i.e., replicate sets of brother-sister mating). To minimise opportunities for researcher applied selection on productivity, one replicate vial with at least 16 emergent flies was randomly selected to provide the parents of the next generation (as in McGuigan et al., 2011); two other vials supplied flies for wing measurement in alternate generations as detailed in the main document Methods. Due to COVID-19 related logistics, inbreeding was paused for one generation after 11 generations of mutation-drift evolution, and a second time 15 generations later (i.e., inbreeding was imposed for 30 generations over a 32-generation period: Table S2). In those pause generations, we were unable to collect virgin flies, and >1 male and female per line were allowed to breed. Inbreeding was re-imposed in the following generation, where pairs may have been full or half siblings, or double first cousins. Logistical constraints also precluded wing image collection in the same two generations, and in two other generations (1 and 3 generations prior to the 1$^{st}$ inbreeding disruption) (Table S2).

Lines with <6 replicate vials with emergent offspring in a generation had a secondary, non-inbred, back-up established from one available replicate vial. This non-inbred back-up (consisting of a breeding population of ~20 full-siblings) was used to resurrect the line if none of the single brother-sister pairs successfully bred. Across the total experiment, most (71%) lines were inbred every generation; for lines that remained extant at the completion of the experiment, this reflected 30 generations of inbreeding.

Of the lines with relaxed inbreeding due to requiring resurrection from a non-inbred backup, 47% subsequently went extinct; the low fitness (viability and fecundity) typically also precluded these lines from contributing wing data after resurrection, and thus to estimates of mutational variance. Most (43%) resurrections occurred in the B panel, where low sample sizes (number of lines with wing size data per generation) led to poor model convergence (see below). Only two C panel lines were resurrected, while in panels A, E and D, respectively, 8, 15 and 28 lines were resurrected from pools of flies containing a mix of full siblings, half-siblings and double first cousins. We calculated the average number of generations of inbreeding per panel per time-block to determine the appropriate value of generation to include in the regression analysis to estimate $V_M$ (Table S2). Overall, considering the 80% of lines that remained extant in Panels A, C, D and E (Table S1), and thus contributed most to the estimates of $V_M$, resurrection reduced inbreeding by <1 generation: the average number of inbreeding generations at experimental generation 32 (with 30 generations of brother-sister mating) was 29.7, 29.9, 29.2 and 29.3 for Panels A, C, D and E, respectively.

**Table S2.** The average number of generations of brother-sister inbreeding across all lines extant within each Panel at each time-block. The experimental generations (Exp. Gens) reflect the breeding generations throughout the experiment, irrespective of whether brother-sister inbreeding was applied. The cumulative inbreeding (Cum. Inbred) relates those experimental generations to the number of generations of brother sister mating expected within each time-block (no inbreeding in experimental generations 12 or 28). The average number of realised generations of brother-sister inbreeding (values presented for each Panel) were calculated considering only the MA lines that contributed wing samples in the relevant generations.

| Time-block | 1st | 2nd | 3rd | 4th | 5th | 6th | 7th |
|---|---|---|---|---|---|---|---|
| Exp. Gens | 1-5 | 6-8,10 | 13-16 | 17-20 | 21-24 | 25-27,29 | 30-32 |
| Cum. Inbred | 1-5 | 6-8,10 | 12-15 | 16-19 | 20-23 | 24-26,27 | 28-30 |
| A | 2.97 | 7.61 | 13.25 | 17.28 | 21.38 | 25.15 | 28.94 |
| B | 2.48 | 6.99 | 12.47 | 16.70 | 20.29 | 24.05 | 28.17 |
| C | 3.01 | 7.77 | 13.49 | 17.49 | 21.54 | 25.41 | 29.21 |
| D | 2.87 | 7.70 | 13.00 | 17.18 | 21.30 | 24.55 | 28.61 |
| E | 2.70 | 7.01 | 12.70 | 16.72 | 20.81 | 24.58 | 28.36 |

### PANEL B LINES

The experiment was designed to sample five ancestral genotypes that were relatively evenly spaced across the (heritable) size distribution of the DsGRP. However, the founding genotype for the 2$^{nd}$ smallest size panel (B: DsGRP20) had unexpectedly low fitness. More than 25% of the lines established for this panel went extinct within the first five generations of the experiment, and only 26 lines (45%) were extant at the conclusion (Table S1). The lower sample size was further exacerbated by low breeding success per line resulting in few (or no) flies available for wing phenotyping each generation. The B panel had, on average, <50% as much data as the other panels (Table S1). This low sample size contributed to poor model convergence, where we were unable to estimate mutational variance or test the hypothesis that among-line variance increased. We therefore do not present any results for Panel B.

### SEX SPECIFIC MUTATIONAL VARIANCES AND INTERSEX MUTATIONAL CORRELATIONS

We fit model 1 to data from each sex separately, where sex-specific estimates of mutational variance were not significantly different from one another (overlapping 90% REML-MVN CI), and conclusions identical for sex-specific and pooled estimates. Models did not converge to estimate the 105 covariance parameters in a model with 7 time-blocks by 2 sexes. Considering only the final time block (where all lines were assayed in two generations), we estimated the inter-sex correlation in panels A and E as 0.98 [0.81, 1.15] and 0.99 [0.93, 1.04], respectively. This is consistent with the ancestral inter-sex correlation, which was not statistically distinct from 1.0 (see above), and indicated mutations had consistent effects on size irrespective of the sex. The lower sample size in panel D resulted in sex-specific models failing to converge, while for panel C the very small, statistically indistinguishable from zero, estimate of mutational variance in each sex precluded estimation of the covariance between these estimates.

**Table S3.** Per panel estimates (and REML-MVN 90% CI below in parentheses) of average size ($\bar{X}$) in generation 1 (reflecting pre-mutation, ancestral, size), the mean-standardised mutational variance ($I_M = \frac{V_M}{\bar{X}^2}$, the expected proportional change in mean size after one generation of unit strength of selection on mutational variation where $V_M$ is reported in Table 1). For Panels A, D and E, the rate of evolution of mean size through fixation of new mutations ($\Delta M$) is presented on the measurement (mm) cscale, and as a proportion ($R_M = \frac{\Delta M}{\bar{X}}$) of the generation 1, or global (experiment wide) panel mean (reported in Table 1). Non-directional (micro-environmental) variation in size over the experiment large (Figure S3) relative to $\Delta M$, suggesting the global mean standardisation is likely the most informative estimate of mean size (and thus $I_M$ and of $R_M$). Nonetheless, mean size estimated in generation 1 differs only slightly from the experiment-long average size of each panel (shown in Table 1), and the estimated $I_M$ (see Table 1 for global mean estimate) and $R_M$are thus also very similar when considered relative to the generation 1 and the global average.

| Panel | $\bar{X}$ (mm) | $I_M$ (x $10^{-6}$) | $\Delta M$[1] (mm x $10^{-4}$) | $R_{M_{Gen1}}$ (x $10^{-2}$) | $R_{M_{global}}$ (x $10^{-2}$) |
|---|---|---|---|---|---|
| A | 1.33 | 1.93 | -2.80 | -2.11 | -2.13 |
| | (1.24, 1.42) | (1.03, 2.89) | (-3.90,-1.73) | (-2.94, -1.30) | (-2.99, -1.31) |
| C | 1.39 | 0.23 | - | - | - |
| | (1.29, 1.48) | (-0.08, 0.54) | | | |
| D | 1.43 | 0.86 | -6.61 | -4.28 | -4.31 |
| | (1.32, 1.53) | (0.34, 1.39) | (-7.17, -5.03) | (-5.12, -3.47) | (-5.16, -3.50) |
| E | 1.46 | 1.82 | -6.36 | -4.37 | -4.45 |
| | (1.35, 1.55) | (0.88, 2.83) | (-7.17, -5.04) | (-5.35, -3.43) | (-5.46, -3.47) |

[1] Consistent with lower 5% CI not including zero, t-tests also supported non-zero slope in each panel. A: $t_{43.8}$ = -4.30, $P$ <0.0001; D: $t_{42.5}$ = -9.32, $P$ <0.0001; E: $t_{44.1}$ = -7.93, $P$ <0.0001.

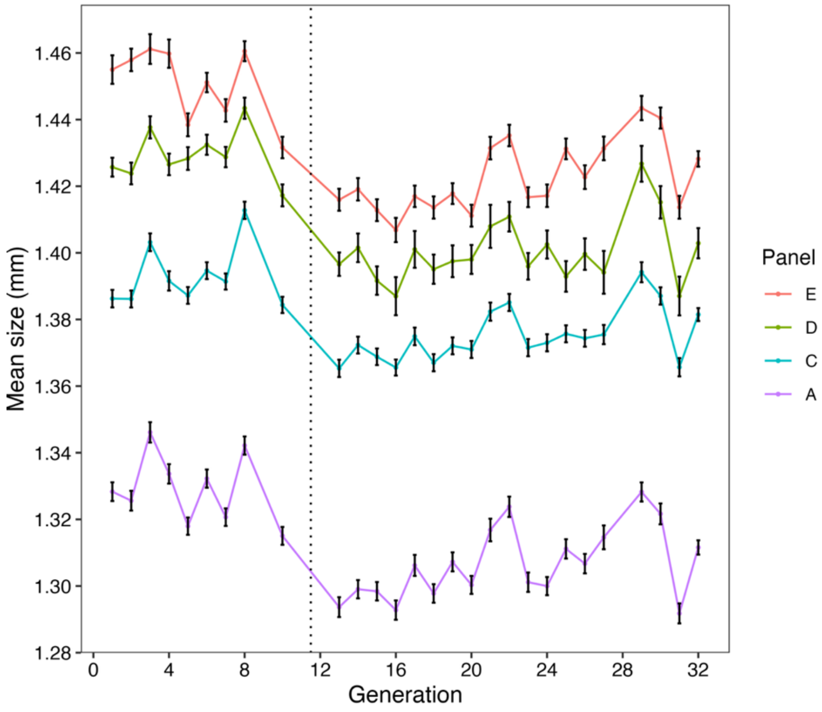


**Figure S3.** Mean size (± SE) of the MA lines in each panel over the experiment. The relatively similar patterns across panels suggest uncontrolled variation in environmental conditions from generation to generation. This micro-environmental variation was accounted for in all models fit to estimate mutational effects. All lines were transferred from one controlled environment facility to another after generation 11 (dotted vertical line). A spike in panel D line extinctions shortly after transfer suggested a difference in conditions, despite identical temperature and day length settings.
